# Labour sharing promotes coexistence in atrazine degrading bacterial communities

**DOI:** 10.1101/581041

**Authors:** Loren Billet, Marion Devers, Nadine Rouard, Fabrice Martin-Laurent, Aymé Spor

## Abstract

Microbial communities exert a pivotal role in the biodegradation of xenobiotics including pesticides^1^. In the case of atrazine, multiple studies have shown that its degradation involved a consortia rather than a single species^2,3,4,5^, but little is known about how interdependency between the species composing the consortium is set up. The Black Queen Hypothesis (BQH) formalized theoretically the conditions leading to the evolution of dependency between species^6^: members of the community called ‘helpers’ provide publicly common goods obtained from the costly degradation of a compound, while others called ‘beneficiaries’ take advantage of the public goods, but lose access to the primary resource through adaptive degrading gene loss. Here, we test whether liquid media supplemented with the herbicide atrazine could support coexistence of bacterial species through BQH mechanisms. We observed the establishment of dependencies between species through atrazine degrading gene loss. Labour sharing between members of the consortium led to coexistence of multiple species on a single resource and improved atrazine degradation potential. Until now, pesticide degradation has not been approached from an evolutionary perspective under the BQH framework. We provide here an evolutionary explanation that might invite researchers to consider microbial consortia, rather than single isolated species, as an optimal strategy for isolation of xenobiotics degraders. Also, we anticipate that future research should focus on the bioaugmentation with stabilized and tightly structured microbial degrading consortia as an effective solution for *in situ* bioremediation of sites polluted with recalcitrant compounds.

## TEXT

Microorganisms in nature usually co-exist as communities whose complexity is under the influence of local environmental conditions and interindividuals interactions. Spatially structured environments, such as biofilms, are more prone to support coexistence of multiple species consuming the same resource because access to the resource and its metabolic by-products is conditioned by the structure of the environment^7,8^. Interactions of various nature between bacterial species that stabilize the community diversity can then arise and persist, but are most likely to disappear when the structure is disrupted^9,10^. In mixed liquid cultures, competition for a single resource is supposed to lead to competitive exclusion^11^. However, if the breaking down of the resource includes both private and public goods, natural selection could favor the emergence of dependency between species, and therefore coexistence, through adaptive gene loss according to the Black Queen Hypothesis (BQH)^6,12^.

Morris *et al.* formulated three key characteristics for a function to follow the BQH. The function allowing consumption of the resource must be *i)* costly, *ii)* essential and *iii)* leaky. Atrazine is one of the most heavily applied herbicide worldwide which is relatively persistent and mobile in soil, and whose degradation products can be traced in soils decades after application^13,14^. Atrazine can be mineralized by microorganisms and is a source of nitrogen. Its entire biodegradation pathway is known (Fig. 1) and can be split into two parts: the upper part consisting in the dechlorination of atrazine and the removal of the two lateral chains from the *s*-triazinic cycle, followed by the lower part consisting in the complete degradation of cyanuric acid. Genes (*atz* and *trz* families) involved in its mineralization are most often located on plasmids or on catabolic cassettes delimited by insertion sequences^15^. Because of the cost of maintaining plasmids, atrazine genes are easily lost after only a couple of generations in culture media without atrazine^16^. However, in nitrogen limited environments, atrazine degrading capacities must be conserved because mineralization of atrazine leads to the essential delivery of nitrogen^17^. Also, exchange of metabolites, such as desisopropylamine, aminoethanol and desethylamine, are produced and released in the environment during atrazine degradation^18^. Therefore, atrazine biodegradation is a good candidate function to test the BQH predictions, and in particular the emergence of dependency between species in a spatially unstructured environment.

**Figure 1.**
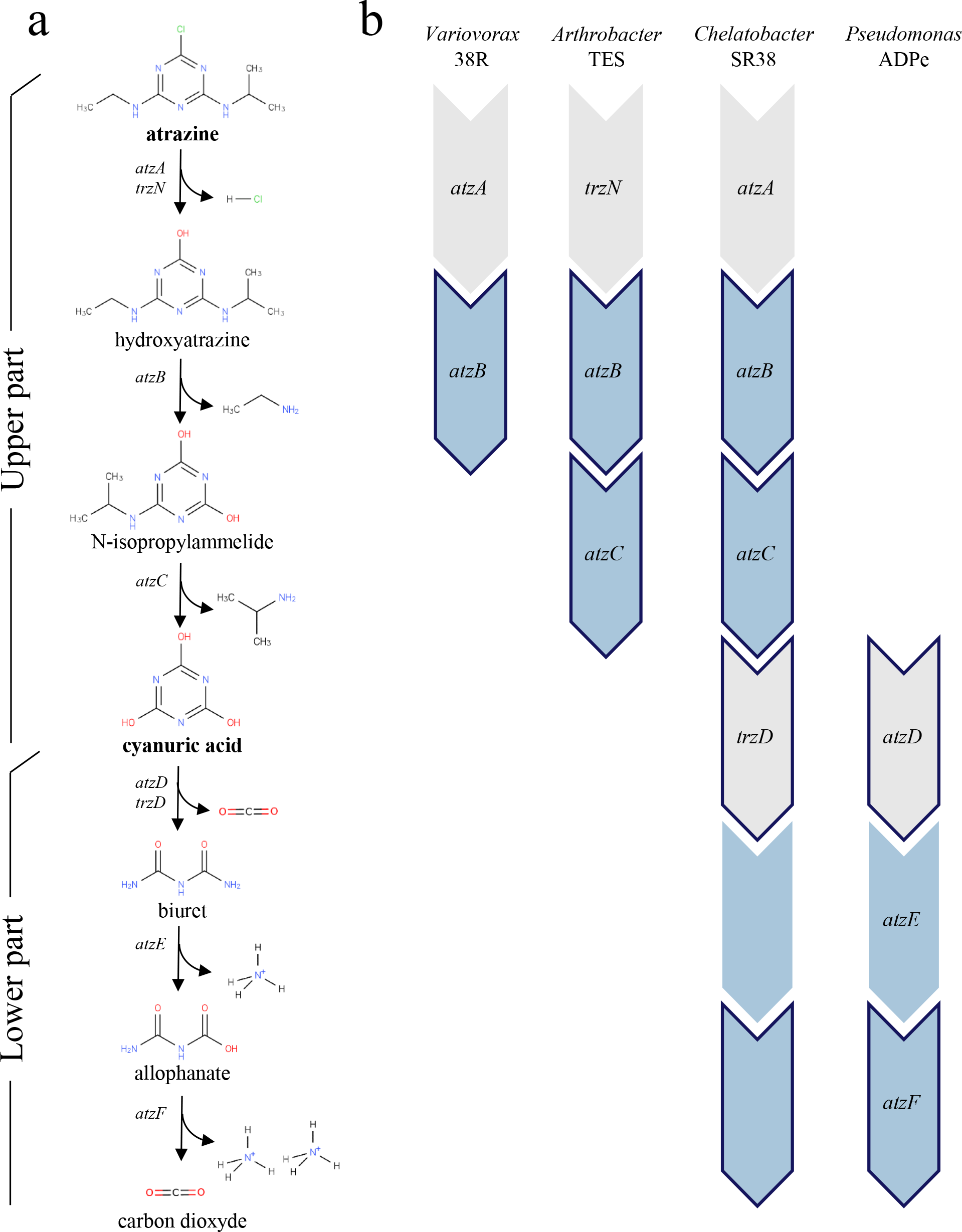
Atrazine metabolic degradation pathway and strains used. **a**, intermediary products of the upper and lower parts of the metabolic degradation of atrazine, as well as genes associated with the different reactions are indicated. **b**, the four ancestral strains, as well as their corresponding atrazine degrading gene repertoire are indicated. Genes encoding enzymes involved in the formation of N-containing by-products are coloured in blue while genes encoding enzymes involved in the formation of C-containing by-products are contoured.

*Chelatobacter* sp. SR38^19^, *Pseudomonas* sp. ADPe^16^, *Arthrobacter* sp. TES^20^ and *Variovorax* sp. 38R^21^ are all originally soil bacteria isolated from arable soils exposed to atrazine. They are all able to at least partly degrade either atrazine or its intermediary products (Fig. 1). They have all been modified to resist to a different combination of two antibiotics in order to evaluate their frequency in mixed culture on double-antibiotics plates. Based on their genetic repertoire for atrazine degradation, it is quite straightforward to predict that in a liquid culture medium with atrazine as the unique nitrogen source, *Arthrobacter* sp. TES and *Pseudomonas* sp. ADPe must coexist because *Arthrobacter* sp. TES will degrade atrazine to supply its nitrogen needs and produce cyanuric acid that will be used by *Pseudomonas* sp. ADPe as its nitrogen reservoir. However, predicting what would happen in a more complex scenario where multiple competing strains evolve in environments containing various nitrogen sources is not trivial. Here, we experimentally questioned whether spatially unstructured atrazine-containing media could support coexistence of multiple atrazine degrading species through BQH mechanisms.

We propagated for ~ 100 generations four-species consortia, in triplicate, in seven minimal media supplemented with citrate in excess, as the carbon source, and either one among three nitrogen sources: atrazine, cyanuric acid: the metabolic product of the upper part of the atrazine degradation pathway, or ammonium sulfate; or 2-way and 3-way combinations of the above mentioned nitrogen sources, keeping the N molarity constant. We found that out of the seven media, all species coexisted in the four ones containing atrazine, *Arthrobacter sp. TES* going extinct in the three atrazine-free media (Fig 2A). The extinction of *Arthrobacter sp. TES* in cyanuric acid supplemented media was expected as it does not possess the metabolic pathway to degrade cyanuric acid, however its extinction also in ammonium sulfate supplemented media is more surprising. Intriguingly, the equilibrium frequencies of the three other species, besides *Arthrobacter* sp. TES, are quite similar after ~100 generations in six out of the seven media, with *Pseudomonas* sp. ADP dominant at ~10^7^ CFU.mL^−1^ and the two remaining species at ~10^6^ CFU.mL^−1^, the exception being the ammonium sulfate – cyanuric acid supplemented medium where *Variovorax* sp. 38R appeared to be dominant at ~10^7^ CFU.mL^−1^.

**Figure 2.**
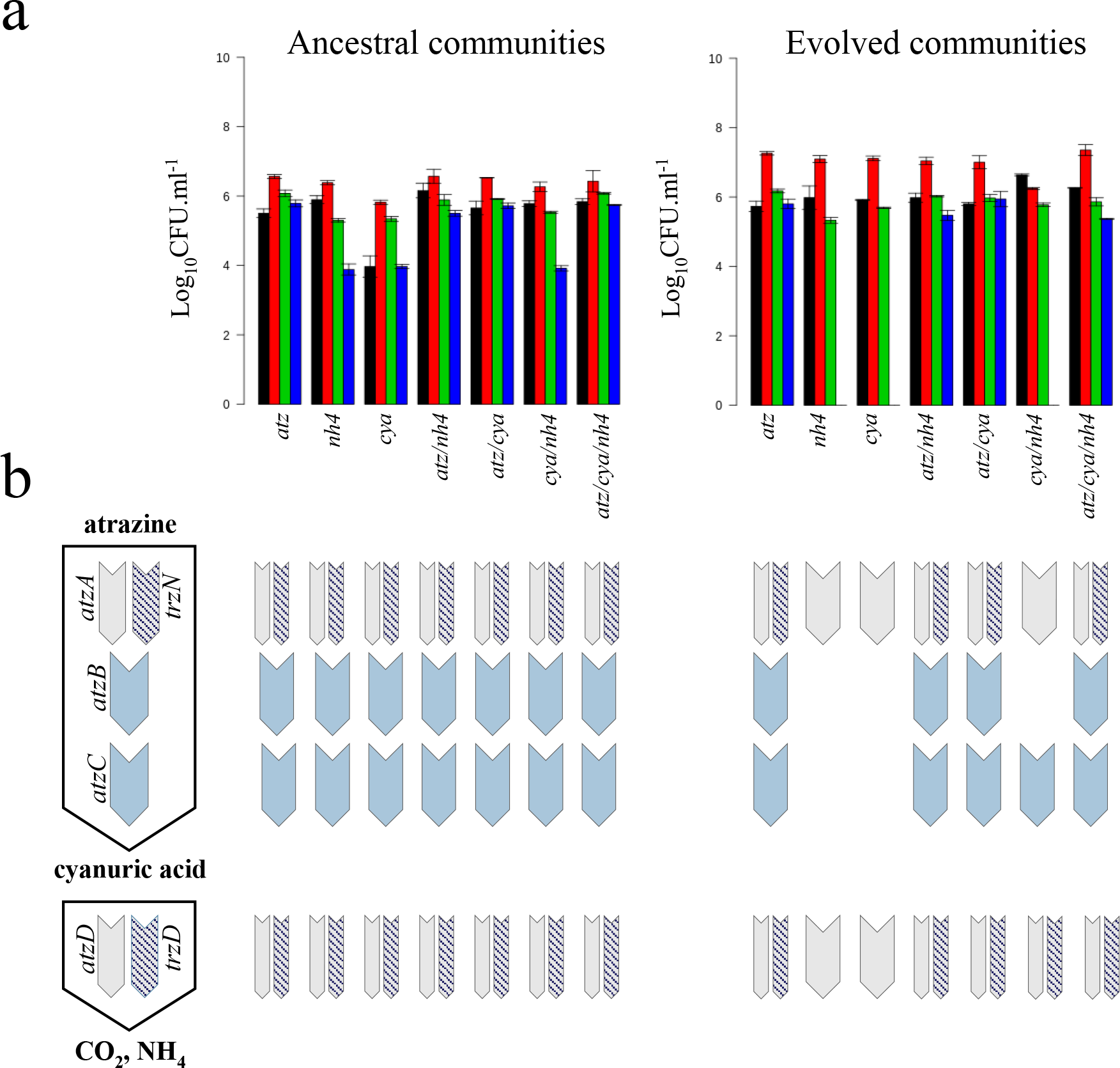
Community composition of the ancestral and evolved consortia across the seven selection media. **a**, species composition of the ancestral and evolved consortia across the seven selection media. Abundances are expressed in log10 CFU.mL^−1^. Black, red, green and blue bars correspond respectively to *Variovorax* sp. 38R*, Pseudomonas sp.* ADPe*, Chelatobacter* sp. SR38 and *Arthrobacter* sp. TES. *atz*, *nh4* and *cya* respectively stands for nitrogen source used either atrazine, ammonium or cyanuric acid. **b**, atrazine-degrading genetic potential of the ancestral and evolved consortia across the seven selection medium evaluated *via* multiplex PCRs. Presence/absence of the corresponding genes is indicated with the presence/absence of the corresponding coloured or hatched rectangle on the model scheme.

We then assessed using multiplex PCRs the presence of the atrazine degradation genes in the evolved consortia across the seven different media. We found that in the four media containing atrazine, all initially present *atz* or *trz* genes were kept in the evolved consortia, preluding the conservation of the atrazine mineralisation potential in these environments (Fig 2B). In the atrazine-free media, only *atzA*, responsible for the dechlorination of atrazine, a step that releases chlorine but does not provide any nitrogen input to cells, and *atzD*, responsible for the first step of the degradation of cyanuric acid, were conserved in the three environments while *atzC* and *trzD* were specifically kept in the ammonium sulfate – cyanuric acid supplemented medium. Since in that specific medium, *atzC* does not confer any advantage to its carrier, *Chelatobacter sp. SR38*, it has likely been carried along because of its colocalization with *trzD* on the same plasmid^22^, responsible for the degradation of cyanuric acid. Rapid evolution, probably through the loss of *atz* genes containing plasmids, therefore led to the loss of the atrazine degradation function in atrazine-free environments, confirming the genetic load of maintaining atrazine genes in atrazine uncontaminated environments.

We then evaluated the atrazine mineralization potential, using ^14^C-atrazine either labelled on the ethylamino chain or on the *s*-triazinic cycle, of the consortia that have evolved in the four atrazine containing environments (Fig 3) and compared them to the ancestral ones. Interestingly, we observed a significant increase of the mineralization potential of the ethylamino chain in evolved consortia compared to ancestors in all but the three nitrogen sources supplemented medium. The observed gain is however much stronger in the atrazine only supplemented medium (from 48 %; CI_95%_=[41.9 – 54.1] to 71.4 %; CI_95%_=[69.4 – 73.4]). When evaluating the mineralization potential of the *s*-triazinic cycle, results are quite contrasted with a significant increase of the mineralization potential in the atrazine only supplemented medium (from 48.3 %; CI_95%_=[36.5 – 60] to 86.9 %; CI_95%_=[81.6 – 92.2]), and significant decreases or no change in the two and three nitrogen sources supplemented media. Altogether, these results indicate that the upper part of the atrazine degradation pathway (*atzA*, *trzN*, *atzB* and *atzC*) has been improved in the evolved consortia, presumably leading to increased intermediary metabolite production, such as aminoethanol, ethylamine or hypoxanthine^18^. While the upper pathway is constitutively expressed, studies have shown that the lower pathway was repressed by the presence of ammonium in the environment^23^. It is therefore quite likely that the stronger accumulation of intermediary metabolites and their consumption by consortium members led to an increased concentration of ammonium in the environment explaining the decreased mineralization potential of the *s*-triazinic cycle in evolved consortia compared to ancestors.

**Figure 3.**
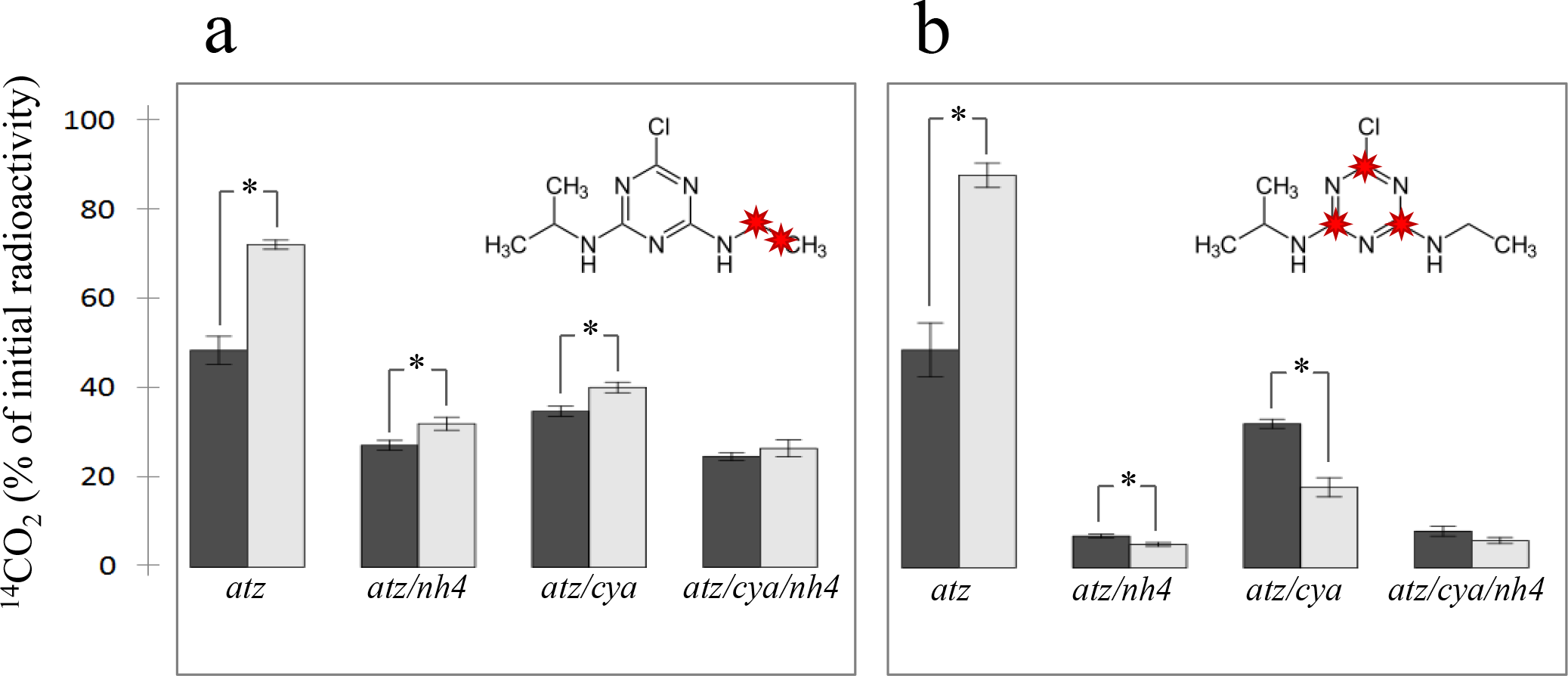
Atrazine mineralization potential of the ancestral and evolved consortia. The mineralization potentials of the ethylamino chain **(a)** and the s-triazinic cycle **(b)** were evaluated with ^14^C-labelled atrazine across the four atrazine-containing selection media. ^14^C-labelled atoms are indicated in red on the atrazine molecule. *atz*, *nh4* and *cya* respectively stands for atrazine, ammonium and cyanuric acid. Significant differences between ancestral (dark grey bars) and evolved (light grey bars) consortia performances are indicated with a star (p < 0.05).

The impressive boost of atrazine mineralization potential observed in the medium supplemented only with atrazine led us to focus on phenotypic and genotypic changes that occurred in this specific environment. We therefore isolated members of the evolved consortia, evaluated individually their genetic repertoire using multiplex PCRs, and assessed their corresponding growth characteristics using Bioscreen^TM^ assays. We found that only *Arthrobacter* sp. TES was able to grow in isolated culture, while the three other members of the evolved consortia depend on *Arthrobacter* sp. TES to grow in this medium (Fig 4A). This acquired dependency for *Chelatobacter* sp. SR38 and *Variovorax* sp. 38R is explained by losses of part of their *atz* genes repertoire (Fig 4B). *Chelatobacter* sp. SR38 lost part of the upper pathway via *atzA* and *atzB* removal, while *Variovorax* sp. 38R kept only *atzA*, responsible for dechlorination of atrazine which does not provide any nitrogen containing by-product. We then wanted to characterize the interactions between members of the evolved consortia. We therefore reconstructed all possible duos and trios, and compared growth performances of each member to their performance in the evolved four-species consortia and in monoculture over 3.5 days (Extended Data Fig 1). A relatively simple and straightforward interaction scheme can be drawn from these results (Fig 4C): *Variovorax* sp. 38R*, Chelatobacter* sp. SR38 and *Pseudomonas* sp. ADP growths are clearly supported by *Arthrobacter* sp. TES. However, *Arthrobacter* sp. TES is penalized in duos or trios with *Chelatobacter* sp. SR38 and *Pseudomonas* sp. ADP, in the absence of *Variovorax* sp. 38R. Interestingly, *Variovorax* sp. 38R by itself has no positive effect on *Arthrobacter* sp. TES, but it counteracts the negative impacts of the two other consortium members on *Arthrobacter* sp. TES. Therefore, coexistence of the four members might be favoured, even though two of the three direct beneficiaries exert antagonistic relationships against the public goods provider, because the third one acts as a buffering intermediary.

**Figure 4.**
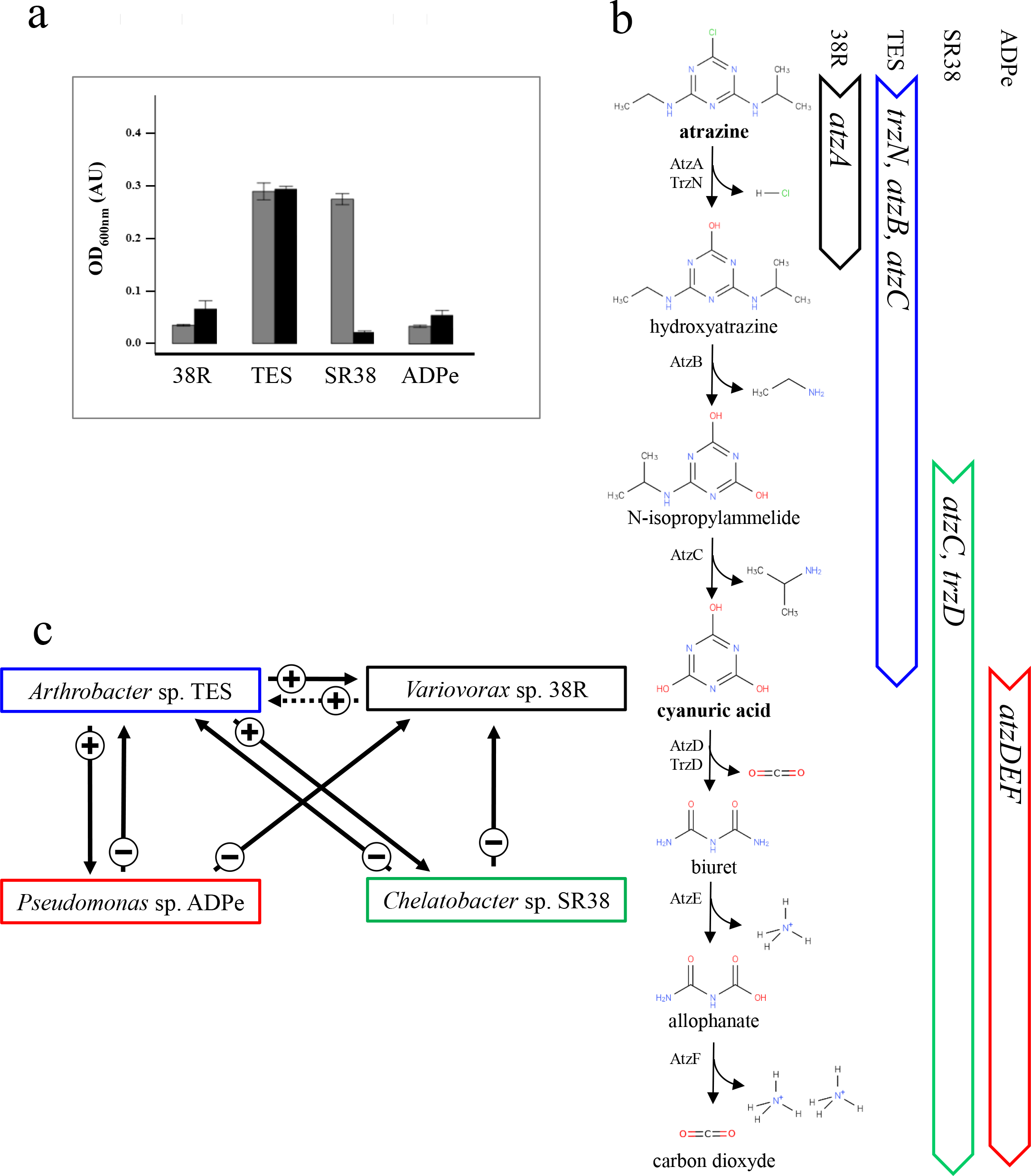
Interspecies dependency for atrazine degradation. **a**, growth performances of the ancestral (grey bars) and community-evolved (black bars) strains estimated as the final OD_600nm_ reached after a 3.5 days cycle. Strains were considered as non-growing if their final OD_600nm_ was < 0.1 Arbitrary Units (AU). **b**, atrazine-degrading genetic repertoire of the community-evolved strains and corresponding metabolic reactions. **c**, interactions scheme drawn from mono- and co-cultures of the four community-evolved strains in the atrazine only supplemented medium. Arrows represent either positive (+) or negative (−) interactions. The dotted arrow expresses the conditional positive effect of *Variovorax* sp. 38R on *Arthrobacter* sp. TES in the presence of the two other strains.

Here, we have witnessed the establishment of dependencies in a four species artificial consortium after evolution in an unstructured, atrazine only supplemented medium during ~100 generations. According to the BQH, dependency was set up *via* adaptive gene loss, and one specific member of the consortium, *Arthrobacter* sp. TES ensuring the costly part of the function, became the provider of public goods to the three other beneficiaries, *Variovorax* sp. 38R, *Chelatobacter* sp. SR38 and *Pseudomonas* sp. ADP. In many cases, pesticide degradation in nature is thought to occur through the involvement of microbial consortia, rather than being the corollary of a single species^4^. Our study provides strong experimental evidences, using the herbicide atrazine as a case study, that the division of labour between populations in microbial communities might be the rule when considering the biodegradation of xenobiotics.

## METHODS

### Experimental Evolution

Four strains with variable atrazine degrading abilities (*Pseudomonas* sp. ADPe, *Chelatobacter* sp. SR38, *Arthrobacter* sp. TES and *Variovorax* sp. 38R, Fig. 1) were cultivated in seven media in four species consortia (n=3) constituting “community” evolution lines. Mineral salt (MS) media (K_2_HPO_4_9.2 mM, KH_2_PO_4_3.0 mM, CaCl_2_0.2 mM, MgSO_4_-7H_2_O 0.8 mM, NaCl 1.7 mM, H_3_BO_3_32.3 µM, FeSO_4_-6 H_2_O 19.2 µM, MnSO_4_-H_2_O 10.6 µM, NaMo 2.1 µM, ZnSO_4_1.2 µM, CuSO_4_ 0.63 µM, biotin 0.4 µM, thiamin 0.15 µM,) containing citrate as carbon source (5 mM) were supplemented with varied nitrogen sources combinations (atrazine, cyanuric acid, (NH_4_)_2_SO_4_, or 2 and 3 sources combinations) with a final nitrogen equimolarity of 1.4 mM (7 media). The evolution experiment was realized in 96-wells 2 mL Deepwell microplate at 28°C without agitation during 23 growing cycles of 3.5 days each. For each new cycle, 35 µL of previous culture was used as inoculum for the next cycle into 700 µL fresh medium (1/20 dilution, (log2(20) × 23 cycles ~ 100 generations)). An aliquot of each cycle culture was mixed with glycerol (30% final concentration) and preserved at −80°C.

### Species composition

We were able to easily identify the four strains in the ancestor and evolved “community” lines because each of them was double-resistant to a different combination of two of the following antibiotics: rifampicin 100mg/L, kanamycin 50mg/L, spectinomycin 100 mg/l, streptomycin 100mg/L. To do so, each bacterial strain was grown in MS medium containing either atrazine or cyanuric acid as the nitrogen source and then spread on solid rich medium (TY) containing the antibiotic for which it was to be resistant. Resistant growing colonies were collected and cultivated in their specific liquid MS medium to ensure that they have kept their metabolic ability towards atrazine. This procedure was repeated to confer a second resistance to each strain. *Pseudomonas* sp. ADPe is Strep^+^/Rif^+^, *Chelatobacter* sp. SR38 is Kan^+^/Strep^+^, *Arthrobacter* sp. TES is Spec^+^/Rif^+^ and *Variovorax* sp. 38R is Spec^+^/Kan^+^.

Ten-fold successive dilutions of T1 and T23 “community” lines were done and inoculated onto LB agar plates supplemented with different antibiotic combinations. After 3-days of growth at 28°C, CFU were counted across 3 dilution levels.

### Atrazine-degrading genetic repertoire

Multiplex PCR targeting *atzA*/*atzD*/*trzD* and *atzC*/*trzN*/*atzB* genes were used to determine the atrazine degradation repertoire of each strain of the ancestral and evolved “community” lines. 3 or 4 isolated colonies from the ancestral and evolved “community” lines were resuspended in water and served as DNA template for PCR reactions. PCR reactions were conducted using the following primers [atzA (5’-TGA AGC GTC CAC ATT ACC-3’, 5’-CCA TGT GAA CCA GAT CCT-3’), atzD (5’-GGG TCT CGA GGA TTT GAT TG-3’, 5‘-TCC CAC CTG ACA TCA CAA AC-3’) trzD5’-CCT CGC GTT CAA GGT CTA CT-3’, 5’-TCG AAG CGA TAA CTG CAT TG-3’] or [trzN (5’-CAC CAG CAC CTG TAC GAA GG-3’, 5’-GAT TCG AAC CAT TCC AAA CG-3’), atzB (5’-CAC CAC TGT GCT GTG GTA GA-3’, 5’-AGG GTG TTG AGG TGG TGA AC-3’), atzC (5’-GTACCATATCACCGTTGCCA-3’, 5‘-GCTCACATGCAGGTACTCCA-3’)] at a final concentration of 10 µM each, the temperature of primers annealing being of 57°C and the elongation time of 1 minute.

### Atrazine mineralisation activity

Ancestors and “community” lines that have evolved on the four media containing atrazine were inoculated at ~0.001 OD_600nm_ in 200 µL fresh medium supplemented with 4000 dpm final of ^14^C atrazine labelled either on the ethylamino chain or on the *s*-triazine cycle in triplicate. Cultures were then placed at 28°C. A whatman^®^ 3mm Chr paper soaked with barite (saturated solution 0.37M) was placed and sealed on the top of the 96-wells plates and replaced periodically during 3.5 days. Dried papers were fluorography printed on Storage PhosphorScreen (Molecular Dynamics^®^) for 2 days. Screens were scanned by a phosphorimager (Storm 860 Molecular Imager). Indirect reading of released radioactivity intensity was done by ImageQuant 5.2 software, and referred to a standard curve. Mineralisation percentages of lines were determinate with respect to initial radioactivity dosage realized on 200 µL of each culture medium with a Beckman^®^ liquid scintillation counter.

### Population dynamics characteristics of the ancestral and evolved line

The population dynamics of ancestral and evolved “community” lines were measured using Bioscreen^TM^. All lines were inoculated at ~0.001 OD_600nm_ in 400 µL of their evolution medium (n=3 for each replicate of each line) in Bioscreen ^TM^ plates. Plates were incubated 3.5 days at 28°C and the evolution of OD_600nm_ was monitiored every 10 minutes after a 10-seconds gentle shake. Population dynamics characteristics of each strain that has evolved in “community” lines was also measured the same way starting from 10 CFU isolated from the evolved “community” lines as inoculums and inoculated at ~0,001 OD_600nm_. Those data were used to determine the maximal OD_600nm_. Cultures that did not reach 0.1 OD_600nm_ at the end of the 3.5 days were considered as non-growing.

### Evaluating interactions between the four evolved species in the atrazine only supplemented medium

For each strain, ten CFU isolated from each community-evolved lines (n=3) in MS-Citrate Atrazine were used as inoculum pools. 4-species consortia, as well as all possible duos and trios were reconstructed and compared to monocultures. Mixtures and monocultures were inoculated at ~0,001 OD_600nm_ and final population densities, after a 3.5 days cycle, were measured for each strain by CFU plating on TY medium supplemented with the appropriate antibiotics combinations.

## ACKNOWLEDGEMENTS

We thank L. Philippot for comments and discussion. Grants from FP7-PEOPLE-2012-IAPP (Industry-Academia Partnerships and Pathways) Marie-Curie project ‘Love-to-Hate’ funded by the European Commission (Grant Agreement number 389 324349) supported this work.

## AUTHOR CONTRIBUTIONS

LB: laboratory experiments, data analysis, manuscript writing. MD: study conception, laboratory experiments, manuscript editing. NR: laboratory experiments. FM-L: study conception, manuscript editing. AS: study conception, data analysis, manuscript writing.

## AUTHOR INFORMATION

The authors declare that no competing interests exist. Correspondence and requests for materials should be addressed to.

## EXTENDED DATA LEGENDS

**Extended Data Figure 1.** Evaluating interactions between the four species evolved in the atrazine only supplemented medium. Growth performances of evolved *Variovorax* sp. 38R (**a**), *Arthrobacter* sp. TES (**b**), *Chelatobacter* sp. SR38 (**c**) and *Pseudomonas* sp. ADPe (**d**) at the end of a 3.5 days cycle in monoculture, as well as in reconstructed duos, trios and in the 4-species consortium were evaluated using CFU plating on TY medium supplemented with the adequate antibiotics combinations. Bars represent mean values with s.e.m (n=3).

