## Supplementary figures and images for "Labour sharing promotes coexistence in atrazine degrading bacterial communities"

### Extended Data Figure 1

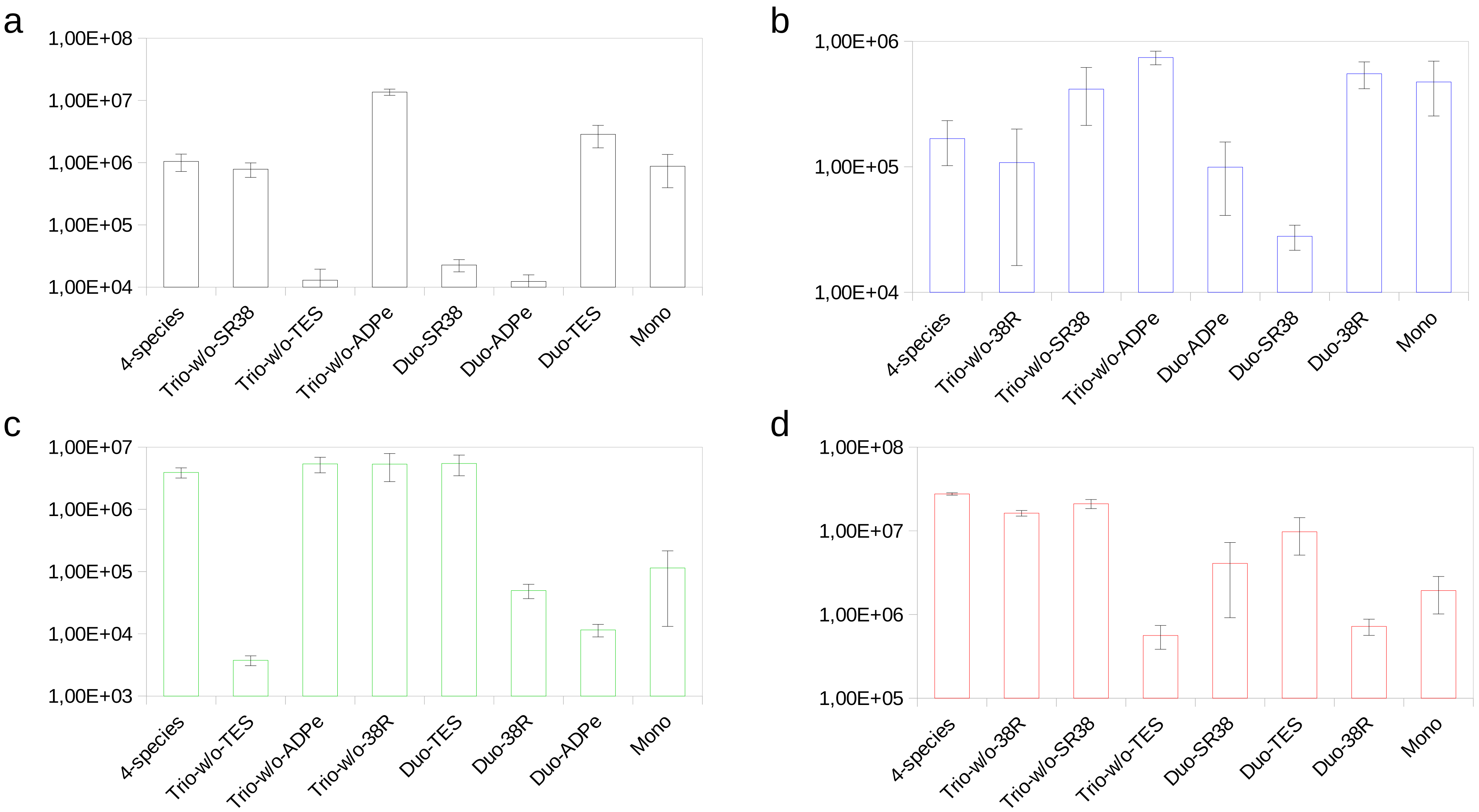
